# Cryo-EM structure of the mechanically activated ion channel OSCA1.2

**DOI:** 10.1101/408716

**Authors:** Sebastian Jojoa-Cruz, Kei Saotome, Swetha E. Murthy, Che Chun (Alex) Tsui, Mark S. P. Sansom, Ardem Patapoutian, Andrew B. Ward

## Introduction

Mechanically activated (MA) ion channels underlie touch, hearing, shear-stress sensing, and response to turgor pressure^1,2^. Previously reported as putative ion channels sensitive to osmolality^3-5^, members of the OSCA/TMEM63 family have been identified as a conserved class of eukaryotic MA ion channels that can be opened directly by membrane tension^6^. The OSCA/TMEM63 family are not homologous to other known MA ion channels, and the structural underpinnings of their function remain unexplored. Here, we report cryo-electron microscopy (cryo-EM) structures of OSCA1.2 from *Arabidopsis thaliana* in lipidic nanodiscs and lauryl maltose neopentyl glycol (LMNG) detergent at overall resolutions of 3.1 and 3.5 Å, respectively. The structures reveal that OSCA1.2 is a trapezoid-shaped homodimer with each subunit containing 11 transmembrane (TM) helices and a folded intracellular domain (ICD) that mediates dimerization. The TM organization of OSCA1.2 has unexpectedly close resemblance to the TMEM16 family of 10 TM helix-containing calcium-dependent ion channels and scramblases^7-9^. We locate the ion permeation pathway within each subunit by demonstrating that a conserved acidic residue is a determinant of channel conductance. Molecular dynamics (MD) simulations provide insights regarding the local membrane environment and lipid interactions, suggesting how OSCA1.2 may be gated by membrane tension. Our work lays the foundation for physiological and biophysical studies on a conserved family of proteins with a newly assigned function as MA ion channels. Moreover, given the role of OSCA proteins as mechanosensors regulating the osmotic stress response in plants^3,6^, our structure could inform strategies to improve plant drought and salt tolerance.

## Main

We purified *Arabidopsis thaliana* OSCA1.2 expressed in HEK293F cells and determined cryo-EM reconstructions in lipidic nanodiscs and LMNG detergent micelle with cholesteryl hemisuccinate (CHS) (Fig. 1a-d, Extended Data Figs. 1,2). The resulting density maps allowed building of 82% of the full-length OSCA1.2 sequence (Extended Data Fig. 3, Extended Data Table 1). Structures of OSCA1.2 in nanodisc and LMNG were nearly identical (RMSD=0.81 Å, Extended Data Fig. 4a-c). We refer to the nanodisc structure throughout unless otherwise noted because of superior resolution and map quality. Importantly, the purified channel reconstituted in liposomes retains MA ion channel activity^6^. Thus, our structures and interpretations correspond to that of a functional OSCA1.2 protein. Additionally, we performed MD simulations at both coarse-grained (CG) and atomistic (AT) levels to model the local membrane interactions of OSCA1.2 (Extended Data Fig. 5).

**Figure 1:**
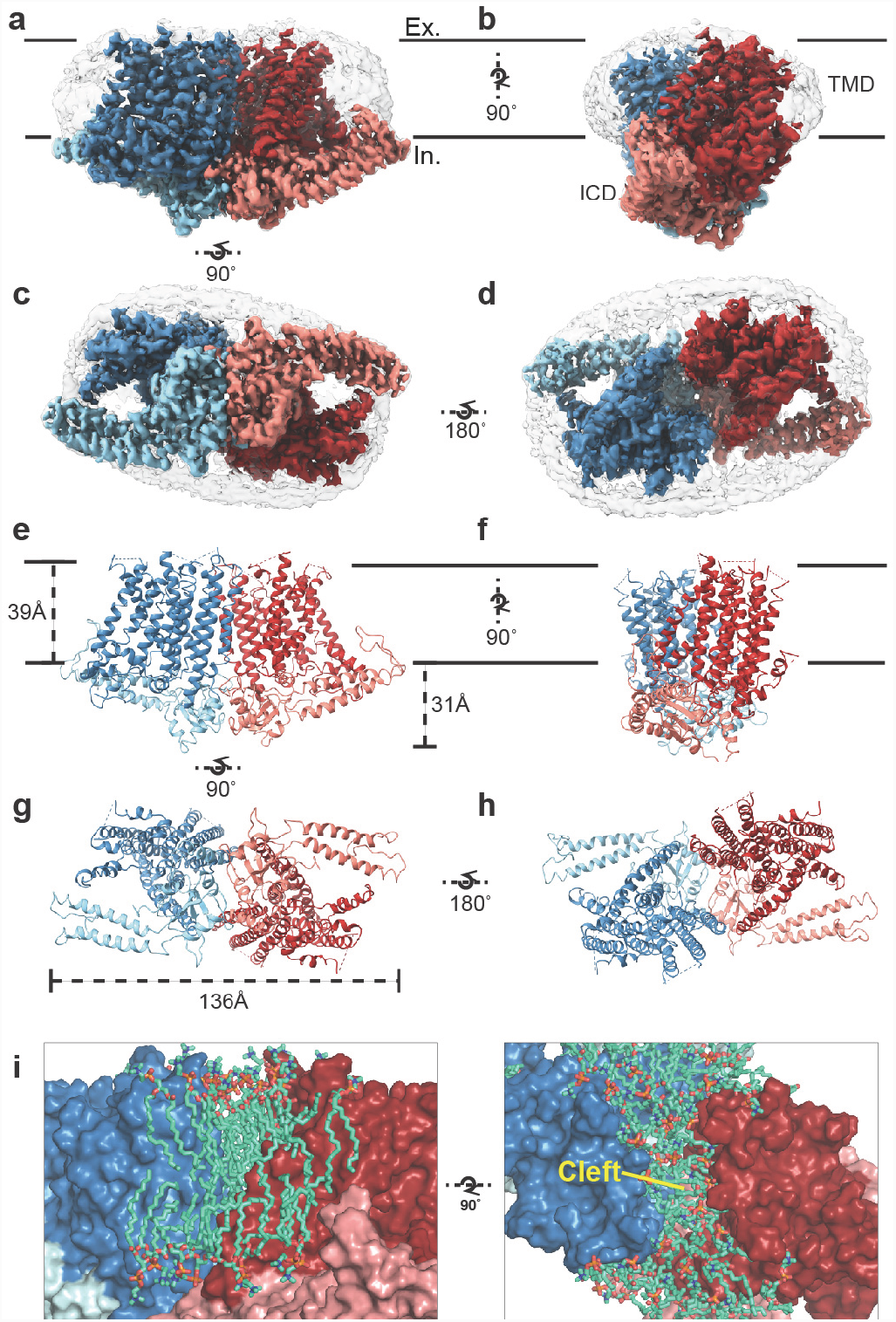
Cryo-EM structure of OSCA1.2 and intersubunit cleft. Side (**a,b**), bottom (**c**) and top (**d**) views of EM density map of nanodisc-embedded OSCA1.2 dimer in nanodisc (sharpened map, blue and red). Nanodisc density is apparent at lower thresholds (unsharpened map, light grey). The intracellular domain (ICD) of each monomer is colored lighter than the transmembrane domain (TMD). Side (**e,f**), bottom (**g**) and top (**h**) views of OSCA1.2 dimer model. Coloring follows the same scheme used in density map. **i**, Molecular dynamics (MD) simulations revealed the inter-subunit cleft to be occupied by phosphatidyl choline (PC) molecules arranged as a bilayer, shown in side (left) and top (right) views (also see Extended Data Fig. 5a) Lipid molecules shown are within 6 Å of residues 367–389 of TM3 (partial), 455–490 of TM5 and 616–685 of TM9–10.

The structure of OSCA1.2 reveals a homodimer with a two-fold symmetry axis perpendicular to the membrane (Fig. 1a-h). The dimer is 139 Å across at its widest dimension. The bulk of the protein lies within the membrane; the transmembrane domain (TMD) of each monomer contains eleven TM helices. Linkers between TM helices and a C-terminal region come together to form an ICD that extends ∼31 Å below the membrane (Fig. 1a-h). ICD contributes the sole dimeric interface in OSCA1.2, with a buried surface area of 1,331 Å^2^ (Extended Data Fig. 4d)^10^, only 4% of the total surface area of the dimer. Within the membrane, inter-subunit contacts are nonexistent. Instead, a cleft with minimum width of approximately 8 Å separates one subunit’s TMD from the other (Fig. 1i). Interestingly, AT-MD simulations of OSCA1.2 in a phospholipid (1-palmitoyl-2-oleoyl-*sn*-glycero-3-phosphocholine, POPC) bilayer, consistently place lipid molecules inside the cleft (Fig. 1i, Extended Data Fig. 5a), indicating a role for lipids in stabilizing the dimeric assembly.

Figure 2a,b show the topology and domain arrangement of an OSCA1.2 subunit. The N-terminus of OSCA1.2 faces the extracellular environment while the C-terminus is intracellular. The majority of the ICD is comprised of the second intracellular linker (IL2), which is over 150 residues and contains a 4-stranded antiparallel ß-sheet (IL2ß1-IL2ß4) with well-conserved sequences across the OSCA family (Supplementary Information), and four helices (IL2H1-IL2H4). Three additional helices, contributed by intracellular linkers 1 (IL1H2) and 4 (IL4H) and the C-terminus (CTH), constitute the rest of the intracellular domain. Other than short linkers, we did not observe structured extracellular domains, presumably due to flexibility.

**Figure 2.**
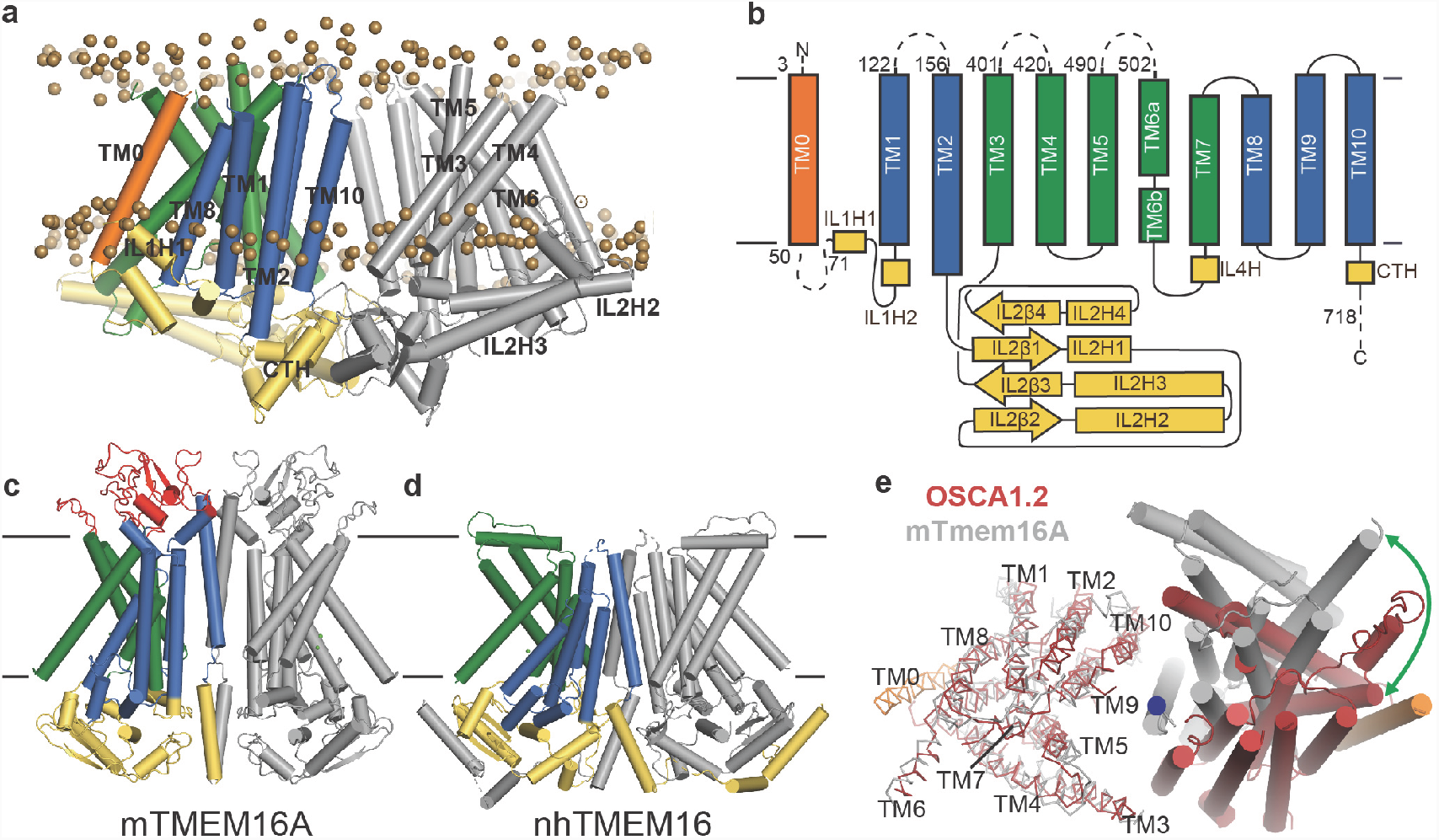
OSCA1.2 topology and comparison to TMEM16. **a**, Structure of OSCA1.2 viewed from the membrane plane. One monomer is colored grey. The other monomer shows intracellular domain in yellow, TM0 in orange, pore lining TM helices in green (TM3–TM7) and TM helices not directly involved in the pore in blue. Phosphorus atoms of POPC molecules within 25 Å of the protein are shown as brown spheres. **b**, Topology diagram illustrating the secondary structure elements of OSCA1.2. Dashed lines represent missing residues in the model. A short coil divides TM6 into TM6a and TM6b. **c**, **d**, Structures of mTMEM16A and nhTMEM16 (PDB: 6BGI and 4WIS, respectively) in the same view as **a**. Extracellular domain of mTMEM16a is colored red. **e**, Bottom view of the transmembrane domains of OSCA1.2 (red) and mTMEM16A (grey) aligned on one subunit (shown as ribbons). The extracellular and intracellular domains have been removed for clarity. The blue circle marks the symmetry axis of OSCA1.2. The green arrow marks the difference in position of TM6 in the non-aligned subunits (cylinders) between the two structures.

As noted previously^4^, a C-terminal region of OSCA1.2 (corresponding to TM4-TM9 in our structure) has loose homology to TMEM16 proteins, which are a family of Ca^2+^-activated ion channels^11-13^ and lipid scramblases^14-16^. Strikingly, our structure reveals that the structural homology extends beyond the C-terminal region; ten of the eleven TM helices of OSCA1.2 closely follow the topology and organization of mouse TMEM16A (mTMEM16A) Ca^2+^-activated chloride channel^8,9,17^ and *Nectria haematococca* TMEM16 (nhTMEM16), which is a Ca^2+^-dependent lipid scramblase^7^ and nonselective ion channel^18^ (Fig. 2c,d, Extended Data Fig. 6a,b). Relative to these TMEM16 structures, OSCA1.2 has an additional N-terminal TM helix positioned at the periphery of the TMD (Fig 2a). To facilitate structural comparison between OSCA and TMEM16 proteins, we refer to this N-terminal TM helix as ‘TM0’, and the remaining TM helices ‘TM1-TM10’. Though TM1-TM10 of OSCA1.2 and TMEM16A align well at the level of a single subunit, the relative orientation of one subunit to the other is different (Fig. 2e). As a result of this distinct dimeric packing, the TMD of the OSCA1.2 dimer is more than 20 Å wider than mTMEM16A or nhTMEM16. Additional features distinguish the overall structure of OSCA1.2 from TMEM16 proteins. First, the intracellular domain of OSCA1.2 is mostly comprised of IL2, which connects TM2 and TM3, while the intracellular domains of mTMEM16a and nhTMEM16 are formed primarily by the N- and C-termini. Second, the regulatory Ca^2+^ binding site composed of acidic and polar residues conserved across the TMEM16 family^7^ does not have a chemical environment conducive for Ca^2+^ binding in OSCA1.2 (Extended Data Fig 6c), consistent with distinct modes of regulation (mechanical activation vs. Ca^2+^ dependence).

In the mTMEM16A dimer, there are two pores; one within each subunit^8,9^. Topological similarities with mTMEM16A (Fig. 2) suggest that OSCA1.2 also has two pores and could conduct ions through a structurally analogous pathway lined by TM3-TM7. The existence of two pores per OSCA1.2 dimer is consistent with the presence of a single subconductance state in stretch-activated single channel currents^6^. To visualize the dimensions of the putative ion permeation pathway, we used the HOLE program^19^ (Fig 3a-c, Extended Data Fig. 7). Towards the extracellular side, this putative pore has an opening greater than 12 Å wide, which narrows into a ‘neck’ approximately 15 Å down the conduction pathway (Fig. 3a-c). The neck extends for over 10 Å and reaches a minimum van der Waals radius of 0.5 Å, closing the channel. Mostly hydrophobic residues (I393, V396, A400, L434, L438, F471, V476, and F515) line the pore’s constriction, as well as several charged and polar residues (Q397, S480, Y468, K512, D523). Interestingly, a pair of π-helical turns (in TM5 and TM6a) are nearby the pore’s neck region (Fig. 3c). Because π-helices are energetically unstable, we speculate that π-to-α transitions at these positions could be associated with gating and, potentially, channel opening. In support of this idea, there are π-helical turns at highly similar locations of TM6 in mTMEM16A^9^ and nhTMEM16^7^, and a π-to-α transition in TM6 underlies Ca^2+^-dependent activation in mTMEM16A^9^ (Extended Data Fig. 6d). Similarly, the pore-lining helices of the epithelial calcium channel TRPV6^20^ undergo π-to-α helix transitions during gating. Directly below the neck, the pore widens, and the side chains of E531, R572, and T568 create a hydrophilic environment that could stabilize hydrated ions (Fig 3c). Further below, the pore widens into membrane-exposed vestibule and is surrounded by IL2 and IL4 before exiting into the cytosol.

**Figure 3:**
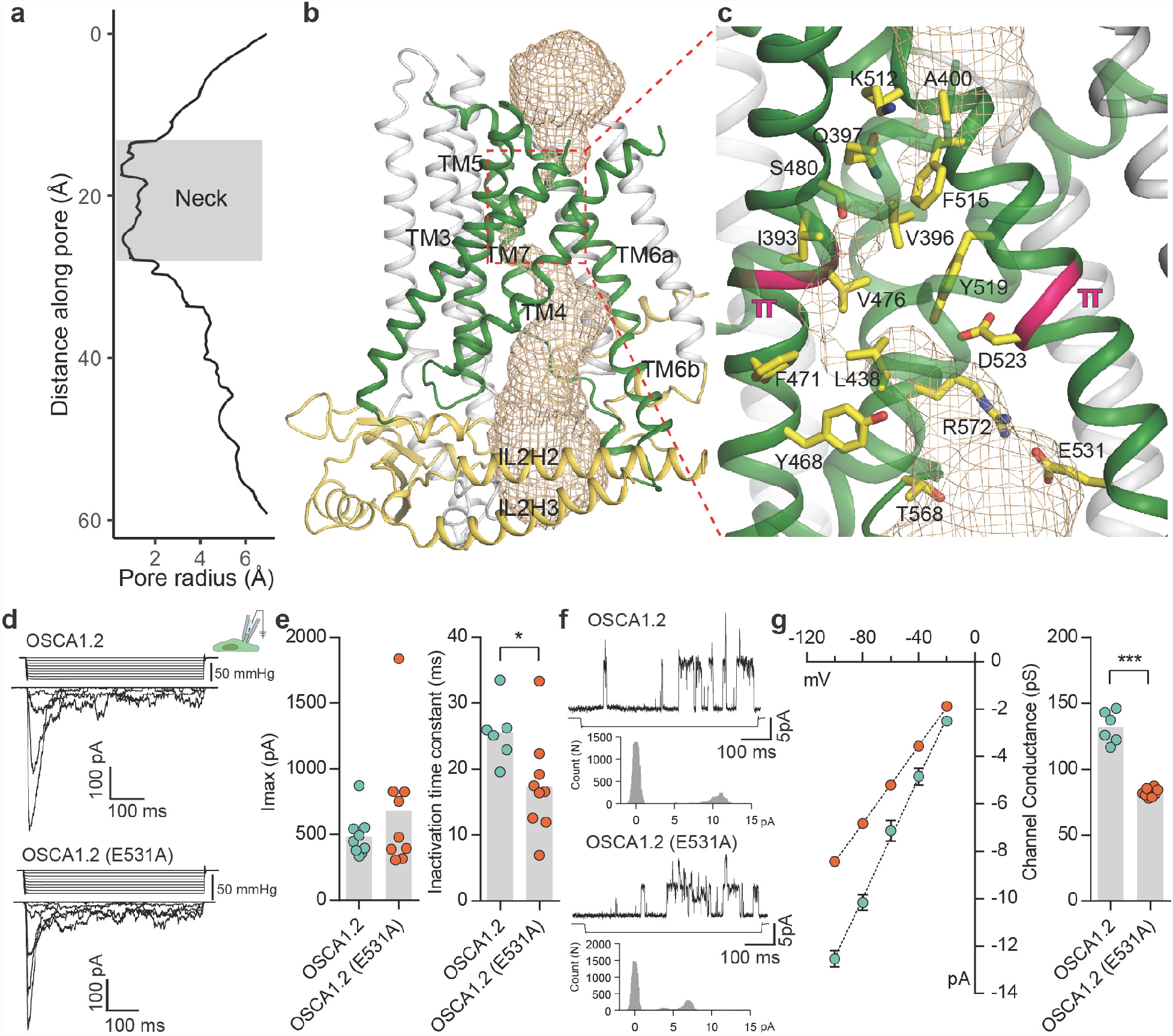
The pore of OSCA1.2. **a**, Van der Waals radii of the pore plotted against axial distance. **b**, Location of the pore in the context of the OSCA1.2 subunit. Pore-lining TMs are colored green, and pore pathway is depicted as wheat mesh. **c**, Expanded view of the neck, and pore lining residues. π-helical turns close to the neck are highlighted pink. **d**, Representative traces of stretch-activated currents recorded from OSCA1.2- or OSCA1.2(E531A)-expressing HEK-P1KO cells. In the cell-attached patch clamp mode, currents were elicited by applying negative pipette pressure in steps of Δ-10 mmHg. The corresponding stimulus trace is illustrated above the current trace. **e**, Left, maximal current response from individual cells expressing OSCA1.2 (N=9) or OSCA1.2(E531A) (N=9). Right, Inactivation time constant (ms) for individual cells across the 2 conditions. Bars represent population mean (OSCA1.2: 25.6 ± 2 ms (N=6); OSCA1.2(E531A): 17.4 ± 2 ms (N=9). \**P*=0.01, Mann Whitney test). **f**, Representative traces of stretch-activated single-channel currents (−80 mV) from wildtype or mutant channels. Channel openings are upward deflections. The negative pressure stimulus is illustrated below the current trace. Amplitude histogram for the representative current trace is represented below the stimulus trace. Single-channel amplitude was determined as the amplitude difference in Gaussian fits of the histogram. **g**, Left, average I-V response curve for stretch-activated single-channel currents from wildtype or mutant channels. Right, Channel conductance from individual cells across the two conditions. Bars represent population mean (OSCA1.2: 132 ± 5 ms (N=6); OSCA1.2(E531A): 82 ± 1 ms (N=8). \*\*\**P*=0.0007, Mann Whitney test).

If TM3-TM7 indeed form the permeation pathway of OSCA1.2, mutation of residues that line the pathway should alter the channel’s permeation or conductance properties. We chose to examine E531, which is the only pore-facing acidic residue conserved in this region across the OSCA/TMEM63 families and thus could contribute to conductance (Fig 3c, Extended data Fig. 6c, Supplementary Information). We recorded and characterized stretch-activated currents in the cell-attached patch clamp mode from cells expressing wildtype or mutant (E531A) channels (Fig. 3d). E531A had maximal current responses comparable to wildtype channels (Fig. 3d,e), and modestly faster inactivation kinetics (Fig. 3e). Strikingly, the mutation decreased the stretch-activated single-channel conductance by 1.6-fold, demonstrating that E531 contributes to the channel’s ion permeation pathway (Fig. 3f,g), and could play a role in binding or sequestering cations. Alternatively, the mutation could alter pore structure and dynamics owing to the interactions of the E531 side chain with R572 and Y605 (Extended Data Fig. 7b). Nonetheless, combining these mutagenesis data with structural homology to mTMEM16A^9,17^, we conclude that TM3-TM7 most likely form the pore. This conclusion is further supported by MD simulations showing hydration of the pore pathway being consistent with the pore radius profile from HOLE (Extended Data Fig. 5b, 7c).

Which structural elements in OSCA1.2 could be involved in membrane tension-sensing and gating? Two notable features present in OSCA1.2, but not TMEM16s, are a hook-shaped loop that enters the membrane that intervenes IL2H2 and IL2H3 (Fig. 2a, 4a), and a horizontal amphipathic helix in IL1 (IL1H1) that slightly deforms the membrane lower leaflet in MD simulations (Fig. 4b). Their association with the cytoplasmic bilayer leaflet suggests these two features could be sensitive to membrane tension, akin to the N-terminal amphipathic helix in MscL^21^, and their movement could be coupled to channel conformation through their interactions with the TMD (Fig 4a,b). Another notable aspect of the OSCA1.2 structure is that the lower region of the pore is exposed to the membrane and thus could permit lipid entry (Fig. 3b, Extended Data Fig. 7a,c). Indeed, in MD simulations, lipids occupy and occlude the cytoplasmic half of the pore, with their phosphate head groups interacting with four lysine residues on TM4 and TM6b (Fig 4c, Extended Data Fig 5c). It is possible that membrane tension promotes gating by affecting the lipid occupancy in this region, analogous to proposed mechanistic models for TRAAK^22^ and MscS^23^. Finally, a general principle of MA ion channels is that their cross-sectional area should increase in response to membrane tension^2,24,25^. Qualitatively, the absence of a dimeric protein-protein interface between membrane domains (Fig. 1i) could allow OSCA1.2 to expand its cross-sectional area significantly in response to physiological lateral tension without the energetic penalty of breaking inter-subunit interactions within the membrane. A comparison of nanodisc-embedded and LMNG-solubilized structures hints at the positional flexibility of the subunits relative to each other, though we would expect larger movements under membrane tension (Extended Data Fig. 4b,c, Supplementary Video 1). The features of OSCA1.2 outlined above might act in concert to modulate channel gating in response to mechanical stimuli (Fig. 4d).

**Figure 4:**
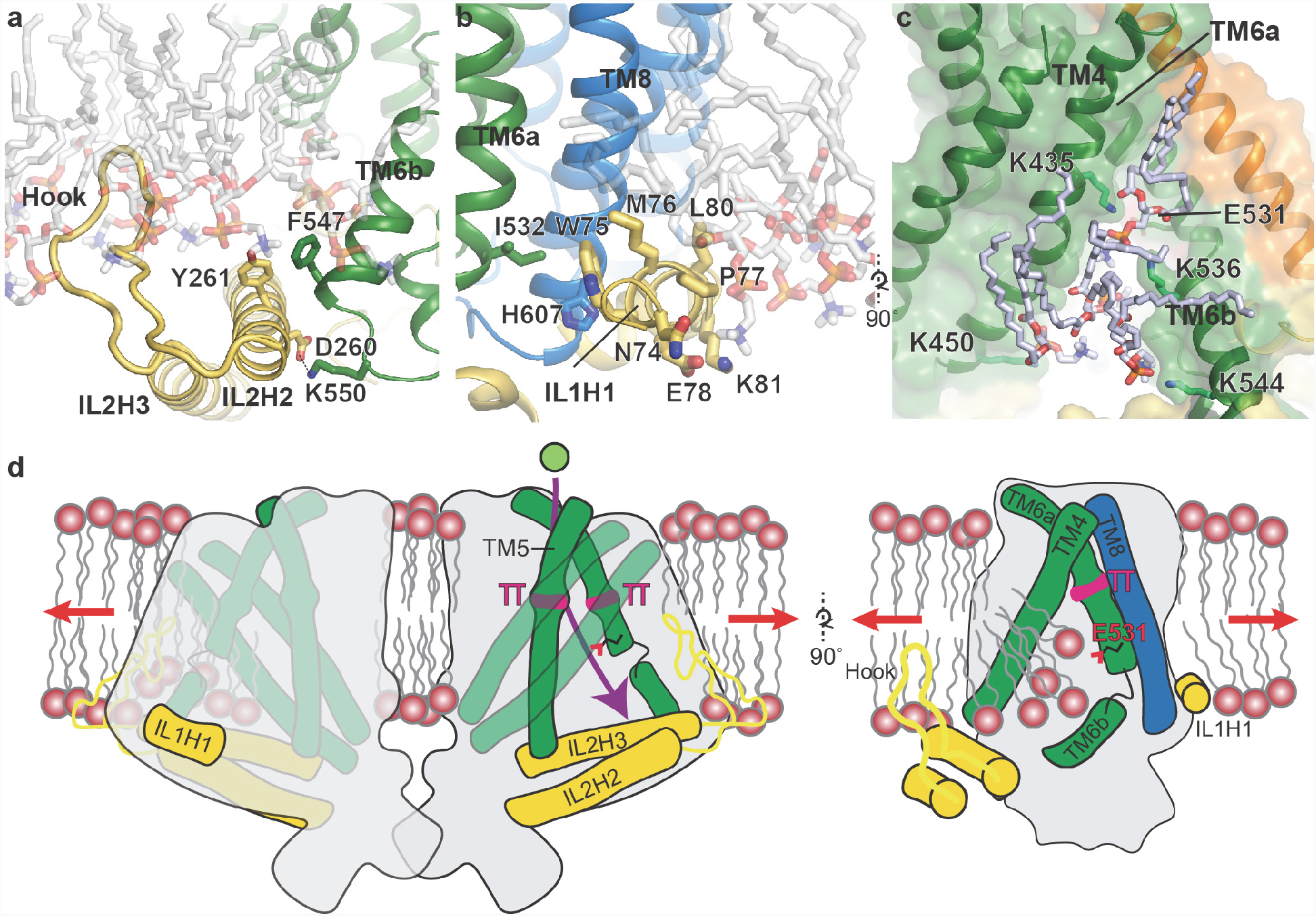
Putative mechanosensitive features of OSCA1.2. **a**, Cartoon representation of the membrane embedded hook, which intervenes IL2H2 and IL2H3. IL2H2 interacts with pore-lining TM6b. **b**, Cartoon and stick representation of the amphipathic membrane-embedded IL1H1, which contacts pore-lining TM6a and TM8. In **a** and **b,** CG-MD snapshots of lipids proximal to the hook and ILH1H1a are shown, demonstrating that both domains are membrane-embedded. **c**, a snapshot (at ∼50 ns) of the MD simulation showing four phospholipids occupying a cytoplasmic cavity of the pore, which is formed by TM4 and TM6b. Interactions of three lipids with the sidechains of K435, K536 and K544 were sustained throughout the 1 µs of CG simulation and also throughout the entire 50 ns of AT simulation (See also Extended Data Fig. 5b). **d**, Schematic showing various features of OSCA1.2 proposed to be involved in MA ion channel function. Red arrows depict direction of membrane tension when applied, though the current structure is not under tension. Green sphere represents ion entering the pore and purple arrow shows pore pathway that transits between π helices. The right panel highlights the amphipathic and membrane embedded regions of one monomer of OSCA1.2 and the local perturbation of the inner leaflet, including entry of lipids into the cytoplasmic side of the pore.

Our structural study of OSCA1.2 elucidates a novel architecture for MA ion channel and reveals unanticipated similarities between OSCA1.2 and TMEM16 proteins, including pore structure. Intriguingly, transmembrane-channel-like (TMC) proteins, which are candidate pore-forming subunits of the hair cell mechanoelectrical transduction (MET) apparatus^26-28^, are distantly related to the TMEM16 family and proposed to have the same topology^29-31^. Although the molecular identity of the MET ion channel remains controversial^27^, the shared TM topology between more distantly related TMEM16s and OSCA1.2 suggests a similar fold for TMCs. It is therefore likely that OSCA/TMEM63, TMEM16, and TMC all utilize a similar architectural scaffold to carry out various functions at the membrane. While the details underlying such functional diversity await discovery, our study establishes a framework to understand how OSCAs exploited this common fold, and evolved specific features, to serve the role of MA ion channels.

## Acknowledgements

We thank W. Anderson for managing the electron microscopy facility at Scripps Research, H. Turner for helping with data collection, and C. Bowman for assistance with computation. We acknowledge J. Kefauver, R. Kirchdoerfer, and members of the Ward lab for helpful advice. K.S. thanks V.K. and D.D. for discussion. This work was supported by a Ray Thomas Edwards Foundation grant to A.B.W, and NINDS grant 1R35NS105067 to A.P. Work in M.S.P.S.’s lab is supported by Wellcome (grant 208361/Z/17/Z), BBSRC (grants BB/N000145/1 and BB/R00126X/1), and EPSRC (grant EP/R004722/1). K.S. is a postdoctoral fellow of the Jane Coffin Childs Memorial Fund for Medical Research. C.C.A.T. is supported by the Skaggs-Oxford Scholarship and the Croucher Foundation. A.P. is an investigator of the Howard Hughes Medical Institute.

## Author Contributions

S.J.C. cloned expression constructs. S.J.C. prepared protein samples and acquired and processed cryo-EM data with assistance from K.S. S.J.C. and K.S. built and refined the atomic structures. S.E.M. carried out electrophysiology experiments and analyzed data. C.C.A.T. performed MD simulations under the supervision of M.S.P.S. A.B.W. and A.P. supervised the project. S.J.C. and K.S. drafted the manuscript, which was edited and finalized with contributions from all authors.

## Methods

### Expression constructs

For structural studies, codon optimized OSCA1.2 gene (UniProt ID: Q5XEZ5) for expression in human cell lines was cloned into vector pcDNA3.1. An EGFP tag was placed at the C terminus and connected to the gene via a PreScission Protease cleavable linker (LEVLFQGP). A FLAG tag (DYKDDDDK) was added to the C terminus of EGFP with two intervening alanines as a linker. We refer to this construct as OSCA1.2-PP-EGFP.

### Protein expression and purification

To obtain samples of OSCA1.2 solubilized in LMNG, four liters of HEK293F cells were grown in Freestyle 293 expression media to a density of 1.2-1.7×10^6^ cells/mL. Each liter was transfected by combining 1 mg/L of the construct with 3 mg/L of PEI in 30 mL of Opti-MEM and then adding the mix to the culture of cells. Transfected cells were grown for 48 hours and then pelleted, washed with ice cold PBS, flash frozen and stored at -80°C for future use. From this point forward, every step of the purification was carried put at 4°C unless otherwise stated. Pellets were thawed on ice, resuspended in 200 mL of solubilization buffer (25 mM tris pH 8.0, 150 mM NaCl, 1% LMNG, 0.1% CHS, 2 μg/mL leupeptin, 2 μg/mL aprotinin, 1 mM phenylmethylsulfonyl fluoride (PMSF), 2 μM pepstatin, 2 mM DTT) and stirred vigorously for 2-3 hours. Subsequently, insoluble material was pelleted via ultracentrifugation for 45 minutes at 92,387 *g* in a Type 70 Ti rotor. Batch binding of the supernatant was performed for 1 hour with 2 mL of GFP nanobody^32^-coupled CNBr-Activated Sepharose 4B resin that had been previously equilibrated with wash buffer (25 mM tris pH 8.0, 150mM NaCl, 0.01% LMNG, 0.001% CHS, 2mM DTT). Protein bound to resin was spun down for 2 minutes at low speed and washed with wash buffer twice and transferred to a gravity flow column. Resin was wash with 10 CV of wash buffer and transferred to a conical tube using 5 to 8 mL of wash buffer, followed by addition of ∼15 µg of PreScission Protease and rotated overnight. Both resin and supernatant were transferred into a gravity flow column and the flow through was collected and concentrated using a 100 kDa MWCO Amicon Ultra centrifugal filter. Concentrated protein was injected onto Shimadzu HPLC and SEC was performed using a Superose 6 Increase column equilibrated to wash buffer. Fractions corresponding to OSCA1.2 were concentrated to 4.4 mg/mL. Typical yields of OSCA1.2 using this procedure were ∼60 µg purified product per liter of HEK293F cells.

To obtain samples of nanodisc-embedded OSCA1.2, the same purification procedures were used with minor changes. Instead of LMNG, solubilization buffer and wash buffer had 1%/0.1% and 0.01%/0.001% n-dodecyl beta-D-maltopyranoside (DDM)/CHS, respectively. After SEC, protein was concentrated down to approximately 200 μL and mixed with MSP2N2 and soybean polar lipid extract (Avanti #541602) at a molar ratio of 1:3:166 (protein monomer:MSP2N2:lipid). Mixture was rocked at 4° C for one hour. 10 mg of Bio-beads SM2 (Bio-rad) prewashed with SEC2 buffer (20 mM Tris pH 8.0, 150 mM NaCl, 1 mM DTT) were added to the mixture and rotated at 4°C. After one hour, another 10 mg of prewashed biobeads were added and rotated overnight at 4°C. Biobeads were removed and protein concentrated and injected onto Shimadzu HPLC for a second SEC using SEC2 buffer and Superose 6 Increase column. Fractions corresponding to OSCA1.2 peak were concentrated to 2.1 mg/mL.

### Cryo-EM sample preparation and data collection

3.5μL of purified protein (at 4.4 mg/mL for LMNG-solubilized OSCA1.2 and 2.1 mg/mL for nanodisc-embedded OSCA1.2) was applied to previously plasma-cleaned UltrAuFoil 1.2/1.3 300 mesh grids and blotted once for 3.5 seconds with blot force 0 after a wait time of 12 seconds. Blotted grids were plunge frozen into nitrogen-cooled liquid ethane using a Vitrobot Mark IV (ThermoFisher). Images for LMNG-solubilized OSCA1.2 were collected at 300 kV using a Titan Krios (ThermoFisher) coupled with a K2 Summit direct electron detector (Gatan) at a nominal magnification of 29,000x with a pixel size of 1.03 Å. 51 frames were collected per movie for a total accumulated dose of ∼60 electrons per Å^2^ using a defocus range of -1.0 to -2.6 µm. For nanodisc-embedded OSCA1.2, a Talos Arctica at 200 kV was used with a K2 Summit direct electron detector at a nominal magnification of 36000x with 1.15 Å pixel size. 54 frames were collected per movie for a total accumulated dose of ∼60 electrons per Å^2^ using a defocus range of -0.4 to -2.2 µm. Automated micrograph collection was performed using Leginon software^33^, collecting 2536 and 1740 total movies for LMNG-solubilized and nanodisc-embedded OSCA1.2, respectively. Movies were aligned and dose-weighted using MotionCor2^34^.

### Cryo-EM image processing

Image assessment and masking of dose-weighted micrographs was performed using EMHP^35^ and CTF values were obtained with Gctf on non-dose-weighted micrographs^36^. A small subset of micrographs at a range of defocus values were used for manual picking and 2D classification in RELION-2.1^37^ to generate templates for the dataset. For LMNG-solubilized OSCA1.2, template picking on 1357 assessed micrographs was done in RELION-2.1 and followed by particle extraction. Particles were imported into cryoSPARC v0.6.5^38^. For downstream image processing, all parameters in cryoSPARC were kept as default unless otherwise stated. 326,398 particles were subjected to *ab-initio* reconstruction requesting 3 classes. Homogeneous refinement with C2 symmetry imposed was performed using the particles and output structure of the best class (low pass filtered to 30 Å) as reference. Particles from the best 2 classes from the initial *ab-initio* reconstruction run were then subjected to a second round of *ab-initio* reconstruction, this time requesting 4 classes. The best 2 classes (134,337 particles) were refined with C2 symmetry using the initial model obtained from the previous refinement, resulting in the final 3.5 Å resolution map. For nanodisc-embedded OSCA1.2, micrographs with a CTF estimation greater than 5 Å were excluded from further processing. Templates for the dataset were generated as previously described, and particles were picked and extracted in RELION-2.1 on 1,211 micrographs. 788,948 template-picked particles were imported into cryoSPARC. One round of 2D classification was performed and best 2D classes were selected (675,536 particles). Two consecutive rounds of *Ab-initio* reconstruction requesting 3 classes were performed, and particles belonging to the best class were taken for further processing. 92,280 particles from the best class from the last round *ab-initio* reconstruction were subjected to homogeneous refinement with C2 symmetry imposed using the final LMNG solubilized map low pass filtered 30 Å as an initial model, resulting in a 3.2 Å map. To improve quality of the map, particles from micrographs with an estimated resolution (calculated by Gctf) worse than 5 Å were excluded from the dataset. The remaining 91,729 particles were re-extracted and imported into cryoSPARC followed by 2D classification. To obtain an initial model based on this dataset, C2 symmetry imposed homogeneous refinement was performed on particles from the best 2D classes (91,729 particles) using the LMNG-solubilized OSCA1.2 structure low pass filtered to 30 Å as the reference model. Subsequently, this model and its particles were used for heterogeneous refinement (3 classes). Best 3D classes were pooled together and refined, resulting in a 3.2 Å map (76,797 particles). Particles for this map were exported from cryoSPARC and per particle CTF was estimated^36^. A first round of masked refinement was performed in RELION-2.1 using the map generated in cryoSPARC (low-pass filtered to 40 Å) as a reference. The output map from this refinement was then used to create a new mask, which was used in the final refinement, resulting in a 3.1 Å resolution map. Using the parameter --solvent_correct_fsc was useful for obtaining a higher resolution structure.

### Model building and refinement

Manual model building of OSCA1.2 was carried out in coot^39^, iterated with real space refinement using phenix^40^ and rosetta^41^. The atomic model was first built and refined against the 3.1 Å resolution map of nanodisc-embedded OSCA1.2 sharpened to a b-factor of -81 Å^2^ automatically determined by RELION-2.1 postprocessing^37^. Registry was initially aided by loose homology of TM4, TM5 and TM6 to mouse TMEM16A^9^ (PDB ID: 5OYB). The refined structure of nanodisc-embedded OSCA1.2 was then docked into the 3.5 Å resolution map of LMNG-solubilized OSCA1.2, sharpened to a b-factor of -170 Å^2^ in cryoSPARC, then manually adjusted and re-refined. In each structure, the model includes residues 3-50, 71-122, 156-401, 420-490, 502-717, totaling 634 of 771 residues in the full-length sequence of OSCA1.2. Structures were validated by MolProbity^42^, EMRinger^43^, and by computing map-to-model FSC plots using phenix.mtriage^44^. Validations were completed with all side chains in the model. However, side chains of residues 268-287 in the membrane hook were truncated to ß-carbon in the deposited models because of relatively poor density in this region. Structural figures were made in PyMOL^45^ or UCSF Chimera^46^. Pore dimensions were calculated using HOLE^19^. Alignment of the amino acid sequences was done using Clustal Omega^47^ and represented using ESPript 3.0^48^.

### Molecular dynamics simulations

Molecular dynamics simulations were performed using GROMACS 5.0.2^49^ (www.gromacs.org) based on the nanodisc-embedded structure. MemProtMD^50^ was used to convert the structure into a coarse-grained (CG) representation using the MARTINI 2.2^51^ force field, which was then embedded in a band of randomly oriented POPC molecules. Water and 0.15 M NaCl were then added to the periodic simulation box. Initial 100 ns simulation with protein backbone beads position restrained permitted lipid self-assembly (force constant 1000 kJ mol^−1^ nm^−2^). The position restraints were then removed for five runs of a 1-µs equilibrium simulation, during which an ElNeDyn elastic network^52^ with a force constant 1000 kJ mol^−1^ nm^−2^ was applied to the protein (lower and upper cut-off were 5 and 9 Å). Protein-lipid contact analysis was performed with a locally written script that employed a 6 Å cut-off, based on the last 800 ns of the simulation. Final contact values were reported as an average of the two subunits in the five runs. We then converted the final frame of one 1 µs CG simulation to atomistic (AT) detail using a fragment-based protocol CG2AT-Align^53^ (force field: OPLS united atom^54^), solvated again in TIP4P water^55^ with 150 mM NaCl. We then performed a 50 ns equilibrium simulation with protein non-hydrogen atoms position restrained (force constant as above). All CG and AT were performed as NPT ensembles held at 310 K (except CG lipid assembly at 323 K) and 1 bar. The time-step for CG and AT simulations were 20 and 2 fs respectively. For the AT simulation, a velocity-rescaling thermostat^56^ with a coupling constant of 0.1 ps, and a semi-isotropic Parrinello-Rahman barostat^R^ with a coupling constant of 1 ps and a compressibility of 4.5×10^−5^ bar^−1^ were used. The LINCS algorithm^57^ was used to constrain all covalent bonds. Electrostatics were modelled using a Particle Mesh Ewald (PME) model^58^ whereas van der Waals’ interactions were modelled using a cut-off scheme, both with cut-off at 1 nm. Simulations figures were created with VMD^59^ and PyMOL^45^. All figures of simulations represent the final snapshot of the AT simulation unless otherwise stated.

### Generation of mutant, cell culture and transfections

The E531A substitution in OSCA1.2 was generated using Q5 Site-Directed Mutagenesis Kit (New England BioLabs) according to the manufacturer’s instruction and confirmed by full-length DNA sequencing. OSCA1.2-PP-EGFP and OSCA1.2(E531A)-PP-EGFP were transfected and tested in Piezo1-knockout (P1KO) HEK293T cells. HEK293T-P1KO cells were generated using CRISPR–Cas9 nuclease genome editing technique as described previously^60^, and were negative for mycoplasma contamination. Cells were grown in DMEM containing 4.5 mg mL^-1^ glucose, 10% fetal bovine serum, 50 U mL^-1^ penicillin and 50 μg mL^-1^ streptomycin. Cells were plated onto 12-mm round glass poly-D-lysine coated coverslips placed in 24-well plates and transfected using Lipofectamine 2000 (Invitrogen) according to the manufacturer’s instruction. All plasmids were transfected at a concentration of 600 ng mL^-1^. Cells were recorded from 24 to 48 h after transfection. The PP-EGFP fusion at the C-terminus of the protein did not affect channel expression or MA current properties.

### Electrophysiology

Stretch-activated currents were recorded in the cell-attached patch-clamp configuration using Axopatch 200B amplifier (Axon Instruments). Currents were filtered at 2 kHz and sampled at 20 kHz. Leak currents before mechanical stimulations were subtracted off-line from the current traces. Membrane patches were stimulated with a 500 ms negative pressure pulse through the recording pipette using Clampex-controlled pressure clamp HSPC-1 device (ALA scientific). Since the single-channel amplitude is independent of the pressure intensity, the most optimal pressure stimulation was used to elicit responses that allowed single-channel amplitude measurements. These stimulation values were largely dependent on the number of channels in a given patch of the recording cell. Single-channel amplitude at a given potential was measured from trace histograms of 5–10 repeated recordings. Histograms were fitted with Gaussian equations using Clampfit 10.6 software. Single-channel slope conductance for each individual cell was calculated from linear regression curve fit to single-channel *I*–*V* plots.

For cell-attached patch-clamp recordings, external solution used to zero the membrane potential consisted of (in mM) 140 KCl, 1 MgCl_2_, 10 glucose and 10 HEPES (pH 7.3 with KOH). Recording pipettes were of 1–3 MΩ resistance when filled with standard solution composed of (in mM) 130 mM NaCl, 5 KCl, 1 CaCl_2_, 1 MgCl_2_, 10 TEA-Cl and 10 HEPES (pH 7.3 with NaOH).

## Data availability

Cryo-EM maps of OSCA1.2 in nanodiscs and LMNG have been deposited to the Electron Microscopy Data Bank under accession codes XXXX and XXXX. Atomic coordinates of OSCA1.2 in nanodiscs and LMNG have been deposited to the PDB under IDs XXXX and XXXX. All other data are available upon request to the corresponding author(s).

---

**Extended Data Figure 1.**
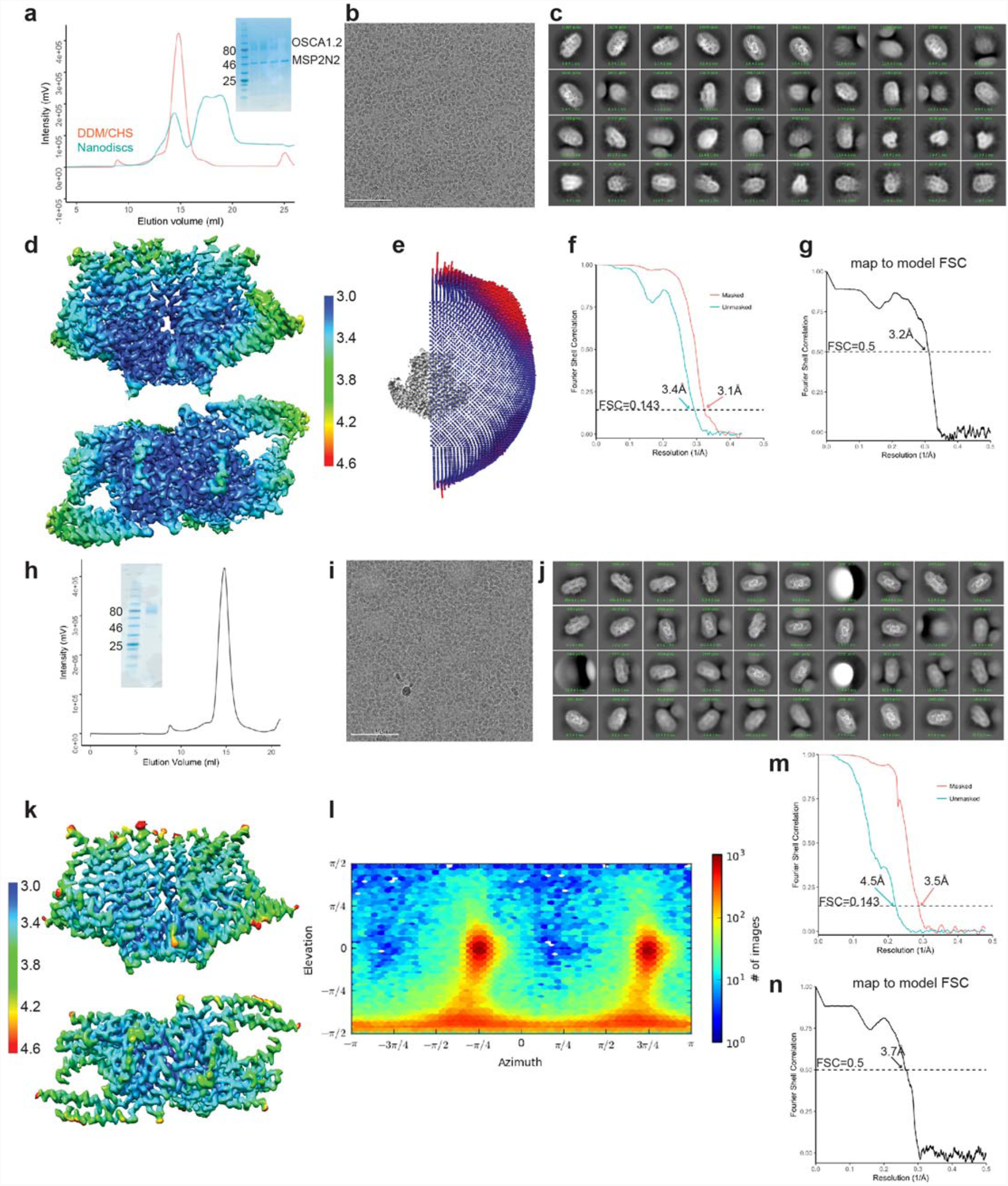
Biochemistry and cryo-EM of OSCA1.2 in nanodiscs and LMNG. **a-g** Correspond to nanodisc-embedded OSCA1.2, and **h-n** correspond to LMNG solubilized OSCA1.2. **a,h**, Representative SEC trace with SDS-PAGE as inset. **b,i**, Representative cryo-EM micrograph. **c,j**, 2D-class averages. **d,k**, Local resolution maps calculated by RELION. Both maps use the same color key, in Å. **e,l,** Angular distribution of reconstructed particles in the refined maps. **f,m**, FSC plots of unmasked and masked maps. **g,n**, FSC of map and model, calculated by phenix.mtriage.

**Extended Data Figure 2.**
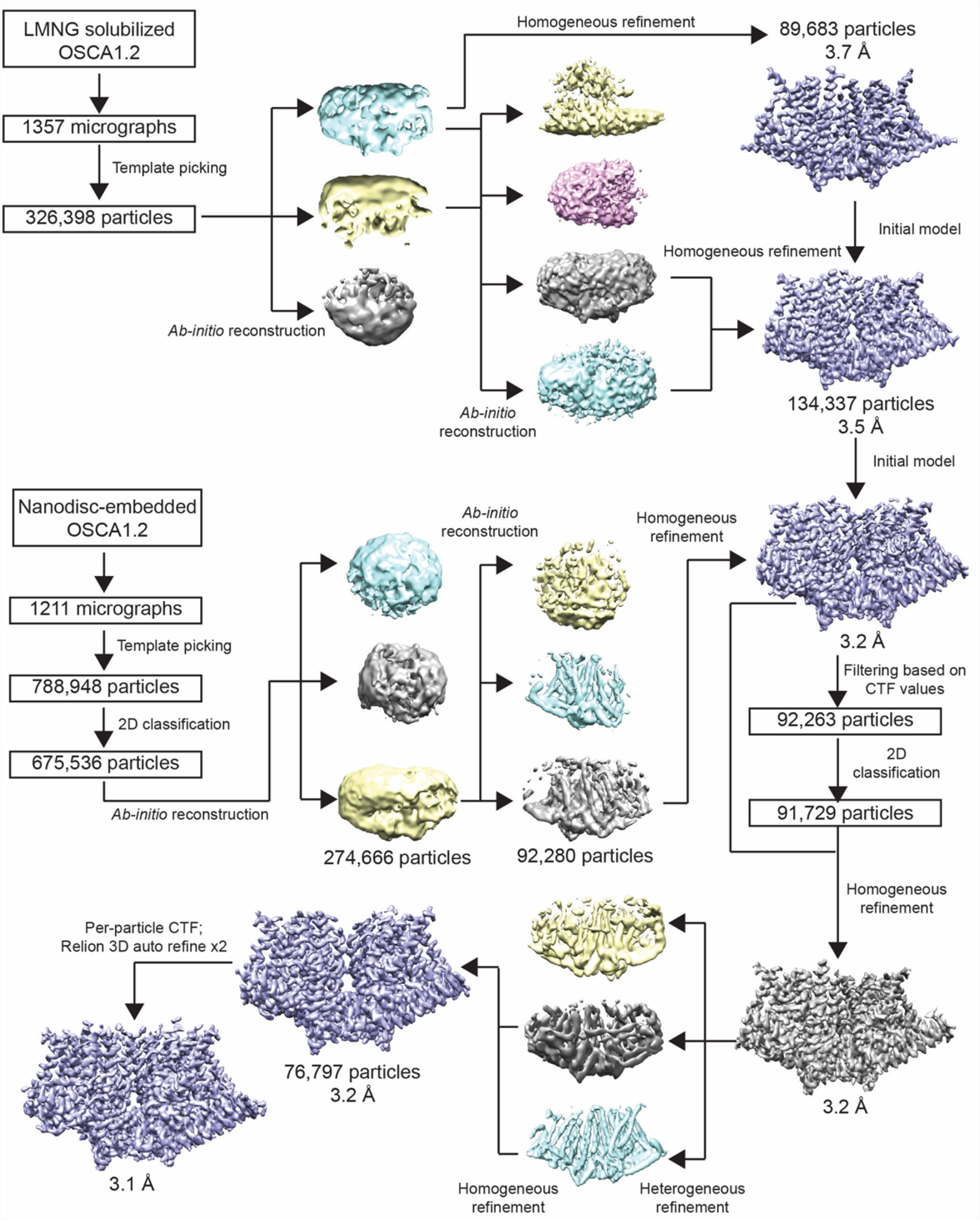
Cryo-EM data processing flowchart. Data processing flowchart for LMNG-solubilized OSCA1.2 map (top) and nanodisc-embedded OSCA1.2 map (bottom).

**Extended Data Figure 3.**
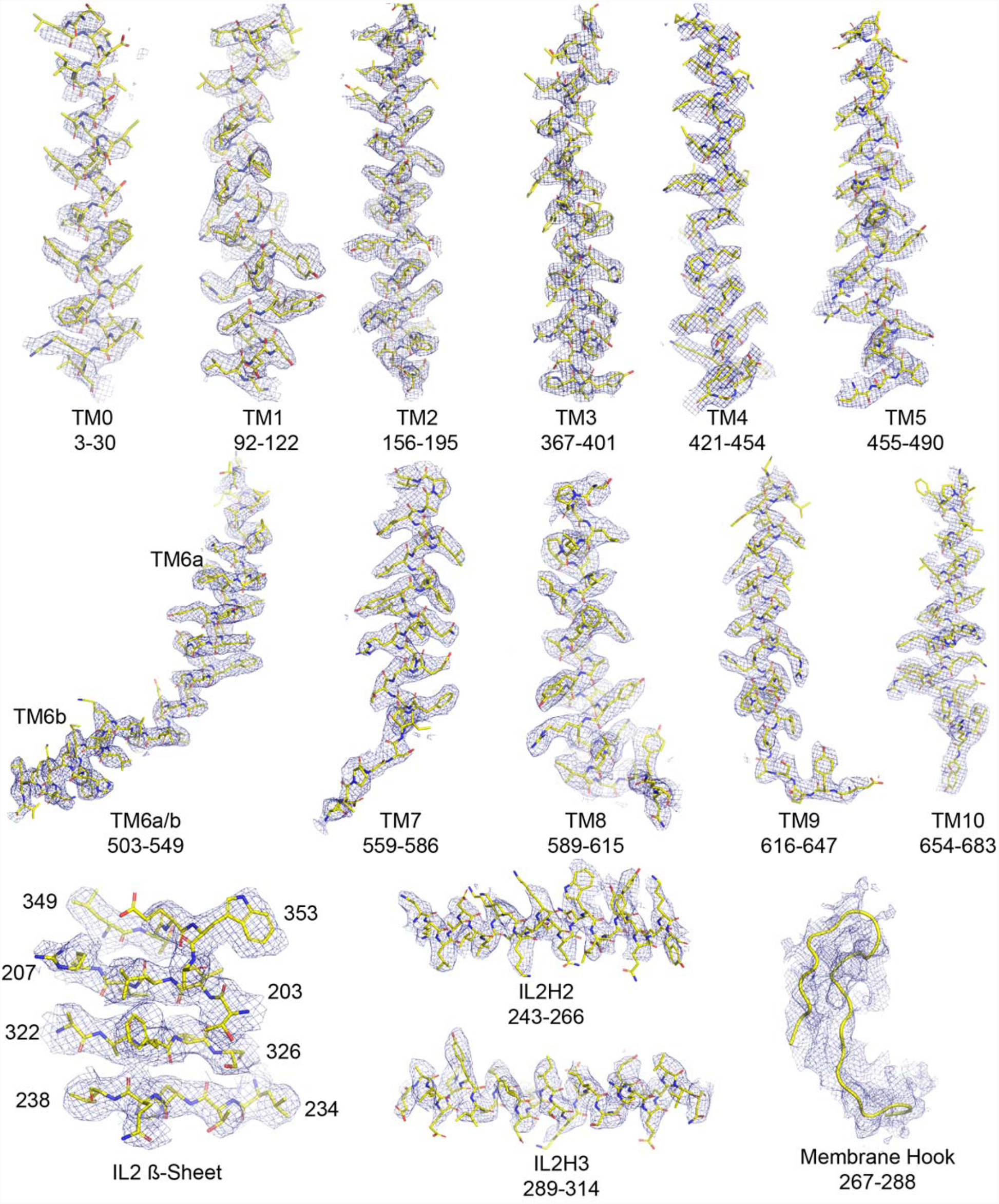
Cryo-EM density and fit to model of OSCA1.2 in nanodisc. Selected regions of cryo-EM map of OSCA1.2 in nanodiscs sharpened with a b factor of -81 Å^2^ superimposed onto the atomic model.

**Extended Data Figure 4.**
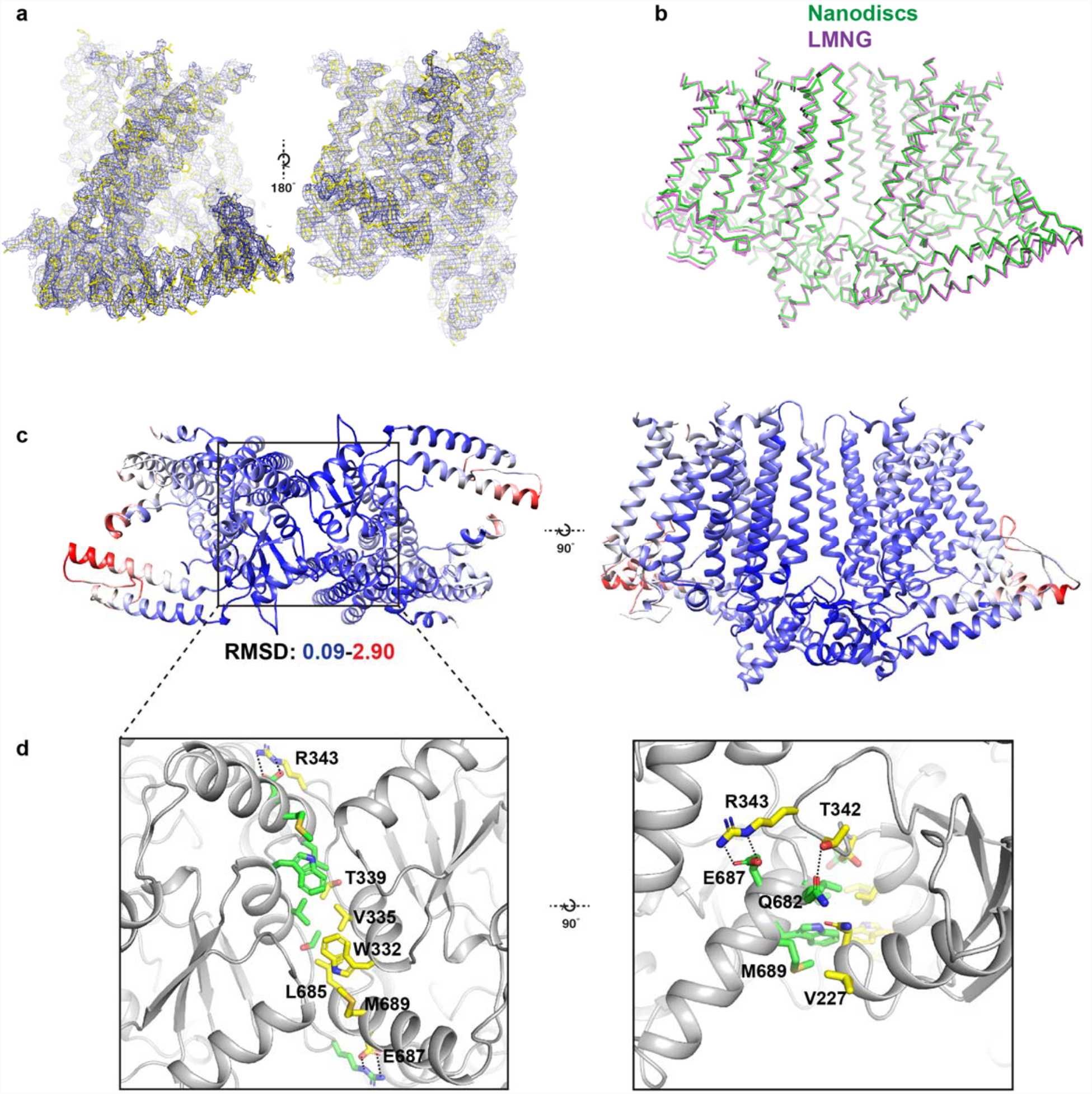
Comparison between detergent and nanodisc-embedded OSCA1.2 and the intracellular dimeric interface. **a**, Cryo-EM map of OSCA1.2 in LMNG sharpened to a b-factor of -170 Å^2^ superimposed into the atomic model. Two side views of the OSCA1.2 monomer are shown. **b**, Ribbon models of OSCA1.2 in LMNG (purple) and nanodiscs (green) aligned and superimposed. **c**, Bottom (left) and side (top) views of the OSCA1.2 model colored by backbone RMSD. Blue color represents RMSD values as low as 0.09 Å, while red values represent values as high as 2.9 Å. White values have intermediate RMSD. Note that RMSD values are lowest at and near the intracellular dimeric interface. **d**, Expanded bottom (left) and side (right) views of the intracellular dimeric interface. Residues at the interface are shown as sticks and colored by subunit. Dashed lines show apparent salt bridges or hydrogen bonds.

**Extended Data Figure 5.**
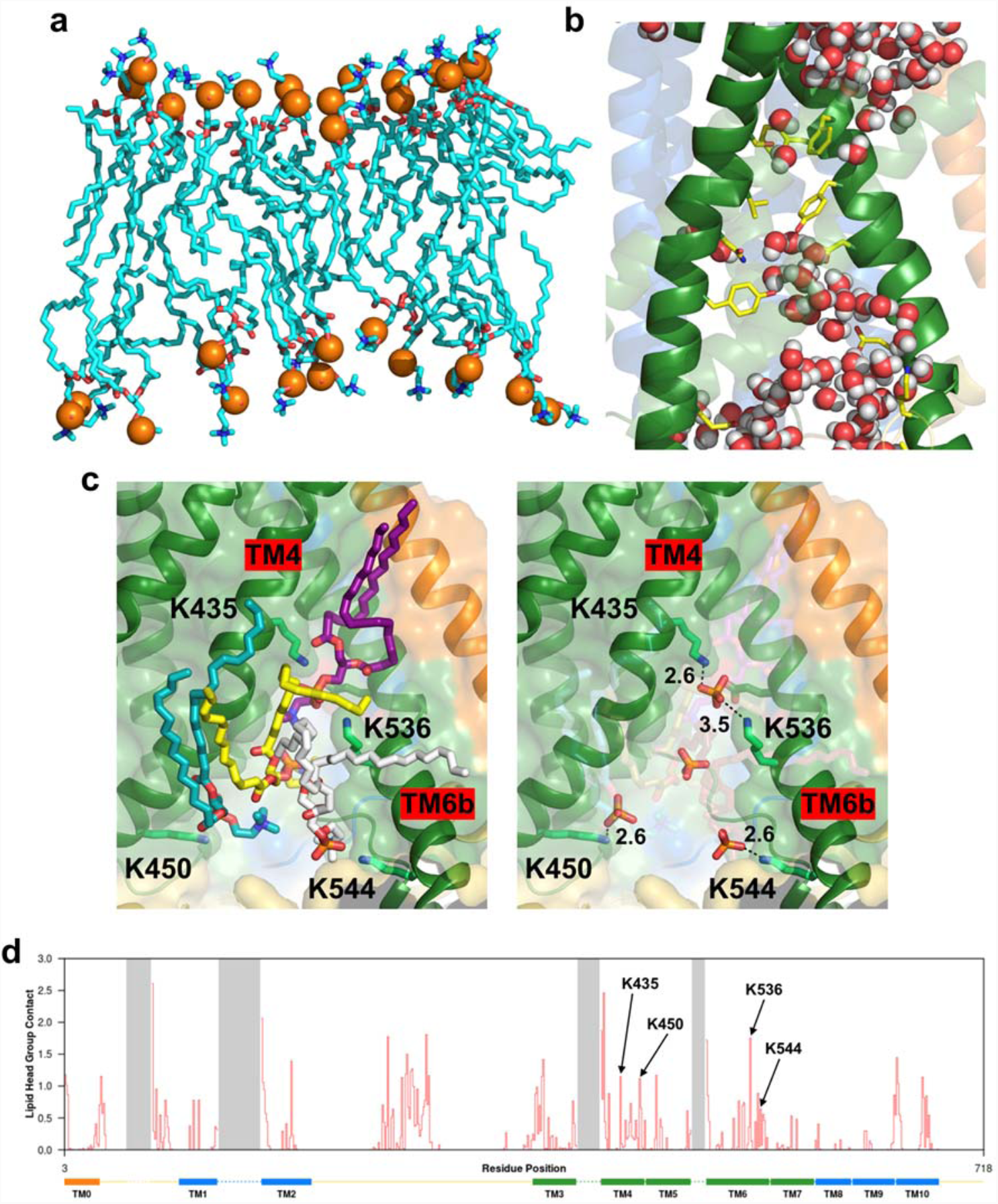
Protein-lipid interactions at the subunit interface and lipid and water molecules within the cytoplasmic region of the pore. **a,** side view of the PC molecules in contact with both subunits at the dimer interface cleft, revealing an asymmetric bilayer between subunits. The outer leaflet of the bilayer in this region contains 18 PC molecules whilst the inner leaflet contains 14 molecules. **b,** Snapshot of an AT-MD simulation at ∼50 ns illustrating the water molecules (red/white) within the two mouths of the pore which are separated by the hydrophobic constriction (compare with Fig. 3c in the main text). **c,** (Left) A snapshot (see also Figure 4c of the main text) highlighting four lipids residing at the cytoplasmic side of the pore taken at ∼50 ns of the AT-MD simulation. (Right) The same snapshot with the lipid phosphates highlighted and the lysine amine nitrogen to phosphate oxygen distance measured in Å, suggesting phosphate head groups are within hydrogen-bonding distance of the lysine residues. **d,** Protein-lipid contact analysis showing the number of phosphate head groups within 6 Å plotted against protein residue number. Transmembrane helices are indicated as rectangles at the bottom and dashed lines represent residues not resolved in our structure. The four lysine residues in **c** are highlighted.

**Extended Data Figure 6.**
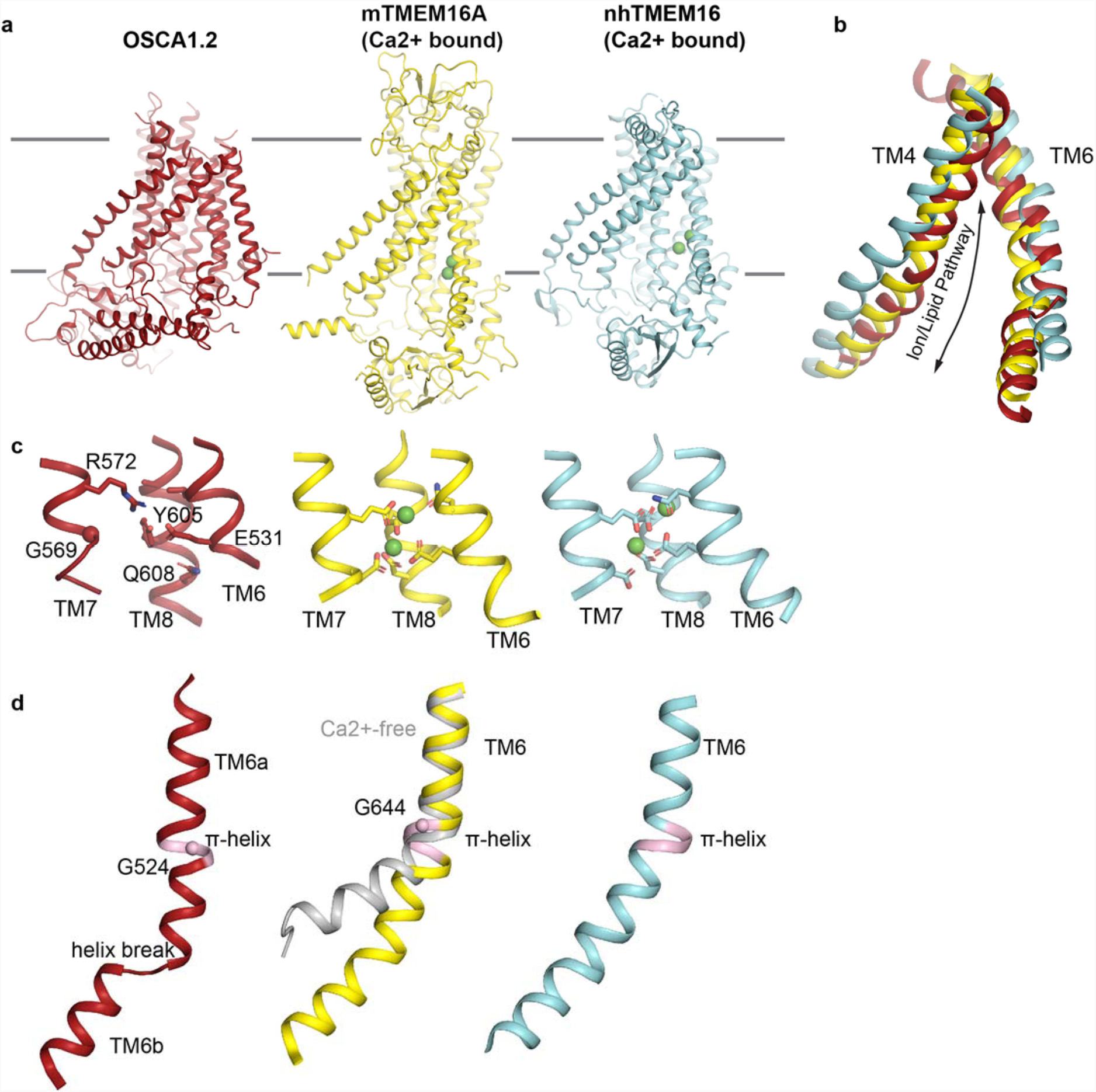
Structural Features of OSCA1.2 and TMEM16 proteins. **a**, Side views of OSCA1.2 (left, red), mTMEM16A (PDB 5OYB, middle, yellow), and nhTMEM16 (PDB 4WIS, right, cyan). Calcium ions are shown as green spheres in mTMEM16A and nhTMEM16. **b**, Superposition of TM4 and TM6 of the structures shown in **a**. The space between TM4 and TM6 in nhTMEM16 is wider than in OSCA1.2 and mTMEM16A, consistent with the lipid scramblase function of nhTMEM16 and the ion channel functions of TMEM16A and OSCA1.2 (ref. 17). **c**, The Ca^2+^ binding site of TMEM16 proteins formed by residues in the cytoplasmic half of TM6, TM7 and TM8 is not conducive for Ca^2+^ binding in OSCA1.2 (left, red). Note E531 in OSCA1.2 is the sole acidic residue in this region. The Ca^2+^ binding sites of mTMEM16A (middle) and nhTMEM16 (right) are also shown. **d**, TM6 in OSCA1.2 (left), Ca^2+^-bound mTMEM16A (middle, yellow) and Ca^2+^-bound nhTMEM16 (right) have π-helical turns (pink). In the Ca^2+^-free structure of mTMEM16A (middle, gray, PDB 5OYG), the π-helix collapses and the helix hinges at a glycine residue of TM6. In OSCA1.2, there is also a glycine at the π-turn in TM6a, raising the possibility of structural similarities in gating transition. The helix break in TM6 of OSCA1.2 could also impart flexibility during mechanical gating.

**Extended Data Figure 7.**
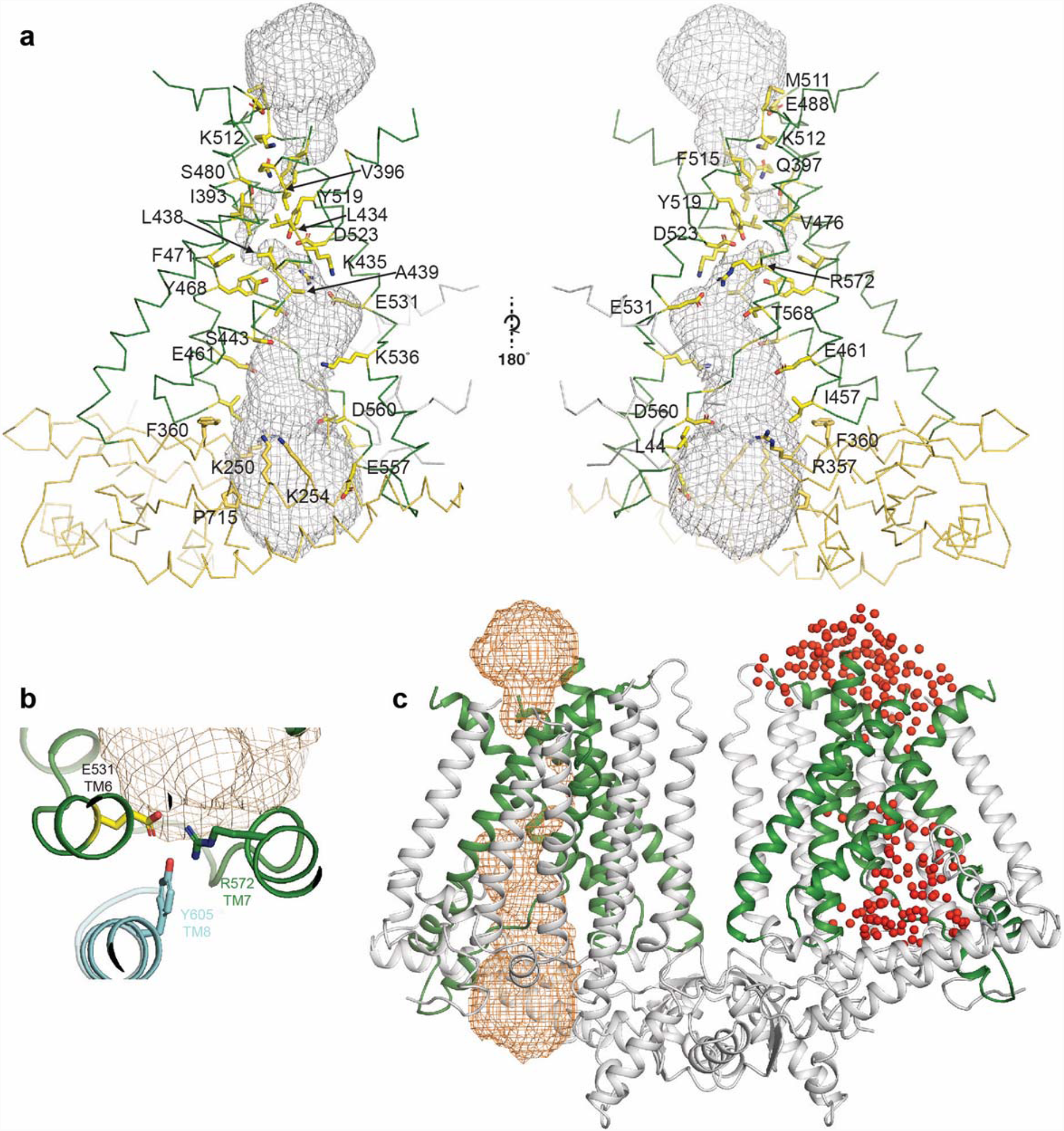
Details of the OSCA 1.2 pore. **a**, Pore and pore lining residues. Side chains are shown as yellow sticks, backbone of OSCA1.2 is shown in ribbon representation and colored green for the TM helices and yellow for the intracellular domain. **b**, Expanded top-down view of E531 and residues that sit in close proximity. **c**, OSCA1.2 dimer showing pore representation obtained using HOLE (ref. 19) in orange mesh (left subunit) and the placement of water molecules (represented as red spheres) around the pore at the end of AT-MD simulation. The similarity between the two supports placement of the pore.

**Extended Data Table 1.** Data collection, processing, model refinement, and validation.

|  | OSCA1.2 in<br>nanodiscs | OSCA1.2 in LMNG<br>detergent |
| --- | --- | --- |
| **Data collection and processing** |  |  |
| Magnification | 36000 | 29000 |
| Voltage (kV) | 200 | 300 |
| Electron exposure (e-/Å<sup>2</sup>) | 60 | 60 |
| Defocus range (µm) | -0.4 to -2.2 | -1.0 to -2.6 |
| Pixel size (Å) | 1.15 | 1.03 |
| Symmetry imposed | C2 | C2 |
| Initial particle images (no.) | 675,536 | 326,398 |
| Final particle images (no.) | 76,797 | 134,337 |
| Map resolution (Å) | 3.1 | 3.5 |
| FSC threshold | 0.143 | 0.143 |
| **Refinement** |  |  |
| Map sharpening *B* factor (Å<sup>2</sup>) | -81 | -170 |
| Model composition |  |  |
| Protein residues | 1268 | 1268 |
| Ligands | 0 | 0 |
| R.m.s. deviations |  |  |
| Bond lengths (Å) | 0.018 | 0.018 |
| Bond angles (°) | 1.57 | 1.65 |
| Validation |  |  |
| MolProbity score | 1.31 | 1.04 |
| Clashscore | 2.89 | 1.3 |
| EMRinger score | 4.45 | 2.73 |
| Poor rotamers (%) | 0 | 0 |
| Ramachandran plot |  |  |
| Favored (%) | 96.5 | 96.9 |
| Allowed (%) | 3.3 | 2.9 |
| Disallowed (%) | 0.2 | 0.2 |

